# Fluid administration rate for uncontrolled intraabdominal hemorrhage in pigs

**DOI:** 10.1101/154203

**Authors:** Ujwal R. Yanala, Jason M. Johanning, Iraklis I. Pipinos, Robin R. High, Gustavo Larsen, William H. Velander, Mark A. Carlson

## Abstract

**Background:** We hypothesized that slow crystalloid resuscitation would improve blood loss and hemoglobin compared to a rapid resuscitation during uncontrolled hemorrhage.

**Methods:** Anesthetized, splenectomized domestic swine underwent hepatic lobar hemitransection. Lactated Ringers was given at 150 or 20 mL/min IV (rapid *vs.* slow, respectively, N = 12 per group; limit of 100 mL/kg). Vital sign monitoring and serum testing was done for up to 180 min, followed by necropsy.

**Results:** Survival was 7 *vs.* 8 out of 12 subjects, rapid vs. slow respectively (p>0.05). The slow group had a less blood loss (1.6 vs. 2.7 L, respectively) and a higher final hemoglobin concentration (6.0 vs. 3.4 g/dL).

**Conclusions:** Using a fixed volume of crystalloid resuscitation in this porcine model of uncontrolled intraabdominal hemorrhage, a slow IV infusion rate produced less blood loss and a higher hemoglobin level compared to rapid infusion.

## INTRODUCTION

Massive hemorrhage and traumatic brain injury each account for about half of early mortality on the modern battlefield.^1, 2^ Preclinical and clinical research directed at these difficult scenarios has evolved management strategies, such as such as rapid evacuation and damage control resuscitation, which have improved outcomes for critically injured personnel.^3, 4^ A central tenant of damage control resuscitation has been the resuscitation of a casualty in hemorrhagic shock with whole blood (preferred) or with blood products using a 1:1:1 ratio of plasma, red blood cells, and platelets.^4, 5^ If blood products are not available, then colloid and/or crystalloid fluids are given in 500 mL boluses “until a palpable radial pulse, improved mental status or systolic BP of 80-90 mmHg is present.”^5^ This latter aspect of damage control resuscitation, in which crystalloid administration is minimized during the transport phase of a subject with hemorrhagic shock until the subject reaches a forward surgical unit, also has been described as hypotensive resuscitation.^4,6, 7^

Avoidance of aggressive fluid resuscitation in trauma victims prior to surgical hemostasis was advocated as early as World War I.^8^ In the modern era, animal^6,9,10^ and clinical^3, 7,11-13^ data have indicated that hypotensive resuscitation improves survival after severe hemorrhagic injury. As a result, the basic tenants of hypotensive resuscitation have been adopted^4^ into the Tactical Combat Casualty Care (TCCC) guidelines^5^ of the United States Army. The current TCCC guidelines^5^ do not explicitly state parameters for either a fluid maximum or an optimal fluid administration rate with respect to the prehospital management of a victim with uncontrolled hemorrhage and shock. The topic of the present study was the issue of optimal fluid administration rate. We hypothesized that for a fixed volume of resuscitation fluid in a porcine model of uncontrolled intraabdominal hemorrhage, a slow rate of intravenous infusion would improve outcomes (blood loss and hemoglobin level) compared to a more rapid rate.

## MATERIALS AND METHODS

### Animal Welfare

Refer to the ARRIVE (Animal Research: Reporting of *In Vivo* Experiments^14^) information in Table S1 of the Supplemental Digital Content. This animal research study was carried out in accordance with recommendations in the *Guide for the Care and Use of Laboratory Animals* (8^th^ ed.) from the National Research Council and the National Institutes of Health,^15^ and also in accordance with the Animal Welfare Act of the United States (U.S. Code 7, Sections 2131 – 2159). The animal protocol was approved by the Institutional Animal Care and Use Committee (IACUC) of the VA Nebraska-Western Iowa Health Care System (protocol number 00760), by the IACUC of the University of Nebraska Medical Center (protocol number 11-064-07-ET), and by the Animal Care and Use Review Office (ACURO) of the United States Army Medical Research and Materiel Command (award number W81XWH-11-1-0836). All procedures were performed in animal facilities approved by the Association for Assessment and Accreditation of Laboratory Animal Care International (AAALAC; www.aaalac.org) and by the Office of Laboratory Animal Welfare of the Public Health Service (http://grants.nih.gov/grants/olaw/olaw.htm). All surgical procedures were performed under isoflurane anesthesia, and all efforts were made to minimize suffering. Euthanasia was performed in accordance with the AVMA Guidelines for the Euthanasia of Animals.^16^

### Study Design and Determination of Subject Numbers

The study design of this report was a non-randomized case-control type (see Discussion). The minimum number of swine (n = 12) utilized in each group was determined with a statistical power analysis^17^ using Δ/σ (Cohen’s *d*, in which Δ is the desired difference in means of numerical data set by the observer, and σ is the estimated standard deviation) = 1.25, false positive rate (α) = 0.05, false negative rate (β) = 0.2, and power (1 – β) = 0.8. The endpoints targeted in the power analysis were blood loss and final hemoglobin level.

### Animal Preparation

Refer to the flow diagram in Fig. 1. Domestic swine (castrated males, 3 months) were purchased from the Agricultural Research and Development Center (Mead, NE) of the University of Nebraska–Lincoln, and acclimatized for 3-5 days under veterinary supervision. Subjects were fed *ad libitum* with corn-soybean meal and water. Subjects were fasted for 12 h prior to surgery, but with no water restriction. Immediately prior to the procedure, animals were premedicated^18^ with a single 3 mL IM injection containing 150 mg Telazol® (tiletamine hydrochloride and zolazepam hydrochloride, 1:1 by weight; Fort Dodge Animal Health, New York, NY), 90 mg ketamine, and 90 mg xylazine.

**Fig. 1.**
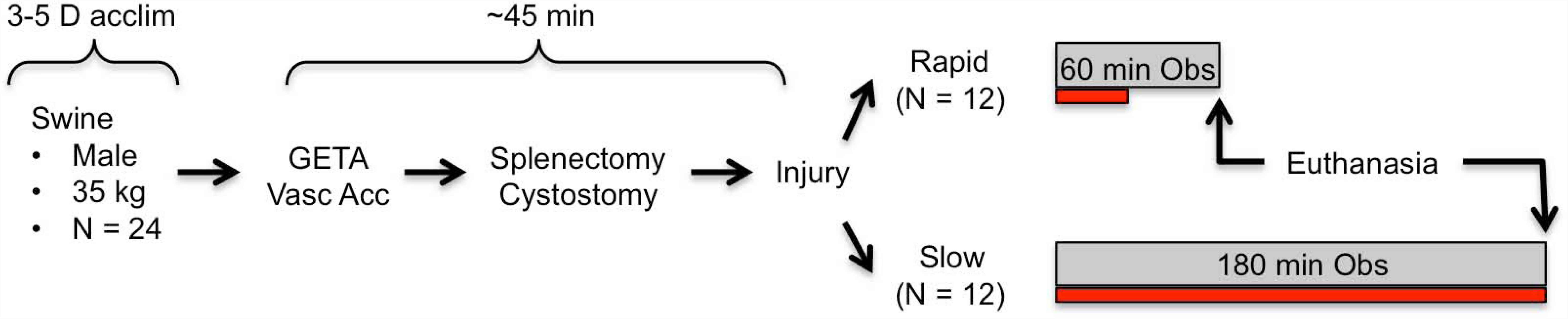
Experiment Flow Chart. Swine were acclimatized (acclim), then on procedure day underwent general endotracheal anesthesia (GETA) followed by access of the carotid and jugular vessels (Vasc Acc), and then other preparatory procedures. After injury, subjects underwent IV crystalloid resuscitation (100 mL/kg maximum volume allowed) for the indicated observation period (Obs). Black bar underneath each observation period indicates relative time required for crystalloid infusion.

Sedated subjects then were weighed, intravenous access was established via an ear vein, endotracheal intubation was performed, and general anesthesia was maintained with 0.5% isoflurane throughout the procedure using a Matrx™ Model 3000 Veterinary Anesthesia Ventilator (Midmark Corp., Versailles, OH). Central arterial and venous lines were placed through a cutdown in the right neck for pressure monitoring, blood sampling and fluid resuscitation. MAP (mean arterial pressure), end-tidal pCO_2_, rectal temperature, cardiac electrical activity, and pulse oximetry were continuously recorded.^19^ Mechanical ventilation was maintained at 12-15 breaths per minute, with a tidal volume of 10-15 mL/kg, in order to keep the end-tidal pCO_2_ at 35-45 mm Hg.^19,20^ Since hypothermia was not an intended variable in this study, a water-circulated warming pad (set at 39°C) was placed under each subject to support body temperature.^21, 22^

A ventral midline laparotomy incision was made, splenectomy was performed,^6, 10, 23-26^ and a transabdominal cystostomy tube was placed.^19^ Per published protocols,^23, 24, 26^ the excised spleen was weighed and then a volume of warm lactated Ringers (LR; 37°C) solution equivalent to three-fold the splenic weight was administered through the jugular line, using a rapid infusion pump (Cole-Palmer Masterflex L/S; Vernon Hills, IL) set at 100 mL/min. Prior to injury, any blood loss incurred during the preparation was quantified by weighing tared surgical sponges that were used to absorb lost blood, and then a volume of LR equivalent to three-fold the pre-injury blood loss (typically <50 mL) was given using the infusion pump.

### Injury Mechanism, Resuscitation, and Observation

Pre-injury vital signs were recorded, the lower half of the midline incision was closed with towel clips, and then the injury mechanism (hepatic left lower lobe hemitransection) was applied, as previously described (a 4 cm cut across the base of the left lateral lobe of the liver^19^), producing a combined portohepatic venous injury. Immediately after injury, the laparotomy incision was closed with towel clips. All the subjects were allowed to bleed without any efforts at local hemostasis (compression, bandage, vessel clamping, etc.).

The goal of the post-injury resuscitation was to give all subjects the same volume of fluid, but with two different administration rates. When the subject’s post-injury MAP dropped below 80% of the pre-injury MAP (defined as the target MAP^19, 25, 27, 28^), LR solution (stored at 37°C^19, 25, 27, 28^) was begun at either 150 or 20 mL/min IV (rapid and slow group, respectively, N = 12 per group) using the infusion pump. The maximum volume of post-injury LR resuscitation was capped at 100 mL/kg.^19^ Resuscitation fluid was administered as long as the MAP was below the target level,^19,20,25,27^ until the animal expired, or until the 100 mL/kg fluid maximum was reached. Similar resuscitation regimens have been utilized in various porcine hemorrhage models by other author groups.^20,25,27,29-31^

The maximum post-injury observation time for the rapid *vs.* slow groups was 60 *vs*. 180 min, respectively. Subjects remained under general anesthesia, with continuous monitoring of vital signs and periodic blood draws for laboratory testing. If a subject was still alive at the end of the prescribed observation period, it was euthanized by increasing the isoflurane fraction to 5% for 5 minutes, followed by bilateral diaphragm incision with transection of the supradiaphragmatic inferior vena cava (intentional exsanguination as approved by the AMVA^16^).

### Endpoints

Heart rate, MAP, pulse oximetry, end-tidal pCO_2_, and rectal temperature were continuously recorded, as described above. Arterial blood samples were drawn pre-injury, at 15 min post-injury, and then at the “final” time point, which was defined as the time of imminent death (for those subjects not surviving the prescribed observation period) or upon completion of the prescribed observation period. Imminent death was defined as MAP ≤20 mm Hg with a pulse pressure of ≤10 mm Hg. Death was defined as MAP ≤20 mm Hg with no identifiable pulse pressure on the monitor’s arterial tracing, end-tidal pCO_2_ <5 mm Hg, and absent corneal reflex.

Immediately after each subject was declared dead, the laparotomy incision was re-opened, and all clots and blood were rapidly evacuated using a combination of tared laparotomy pads, suction, and manual extraction. Tared buckets containing evacuated blood were weighed in order to calculate blood loss. Necropsy was performed immediately after expiration; the liver was explanted for inspection, dissection, photography, and documentation of the injury anatomy.

The complete blood count, prothrombin time (PT), activated partial thromboplastin time (APTT), international normalized ratio (INR), quantitative fibrinogen assay (QFA), and arterial blood gas (ABG) testing were contracted to the Clinical Laboratory of the VA Nebraska-Western Iowa Health Care System. This laboratory used the von Clauss method for the QFA.

### Statistical Analysis

Numerical data were reported as the mean ± standard deviation (SD). For a complete description of the statistical analysis, see the Supplemental Digital Content. Unpaired continuous data were compared with ANOVA; significant results (p < 0.05) were confirmed with nonparametric Kruskal-Wallis analysis of variance. Groups of categorical data were compared with the Fisher exact test. If data for a given subject at a given time point were “missing” (e.g., no data captured secondary to lost blood tube, clotting of a blood specimen prior to running an assay, monitor malfunction, other miscellaneous events), then the respective cells in the data spreadsheet were kept empty (see Statistical Analysis in the Supplemental Digital Content). If a coagulation test was reported as “failed” (meaning, no clot formation during the assay), then the respective data cells were filled as follows: QFA = 20 mg/dL; PT = 37 s; INR = 5 (critical values from contracting laboratory).

## RESULTS

### Pre-injury Data

All raw data from this study and a detailed statistical analysis thereof are contained in the Supplemental Digital Content. Mean subject weights were 35.8 ± 1.8 and 37.9 ± 4.9 kg, rapid *vs.* slow groups, respectively (p > 0.05, unpaired t-test). Pre-injury blood loss (incurred during subject preparation, and including splenic mass) was not different between the two groups (Table 1); splenic mass was 360 ± 71 vs. 343 ± 75 g, rapid vs. slow groups, respectively (p > 0.05, unpaired t-test). Pre-injury fluid administration, which included splenic replacement per the 3x formula described in the Methods, also was not different between the two groups (Table 1). The pre-injury MAP, heart rate, temperature, hemoglobin concentration, platelet count, QFA, PT, and ABG data were not different between two groups (Tables 2-4; see also full statistical analysis in the Supplemental Digital Content). There was a small but significant difference in the INR values between the two groups (Table 4), but this did not seem to be physiologically relevant.

**Table 1.** Blood loss, fluid input, liver weight, and lacerated veins.

| Infusion Rate | Blood Loss (mL)* |  | Fluid input (mL)* |  | Liver weight (g)* | No. Veins Lacerated** |  |
| --- | --- | --- | --- | --- | --- | --- | --- |
|  | Pre-injury | Post-injury | Pre-injury | Post-injury |  | Hepatic | Portal |
| Rapid | 430 ± 151 | 2738 ± 693 | 1288 ± 458 | 3644 ± 523 | 852.9 ± 114.8 | 0.9 ± 0.3 | 1.3 ± 0.4 |
| Slow | 414 ± 48 | 1600 ± 360 | 1138 ± 181 | 3082 ± 1720 | 941.0 ± 127.3 | 1.0 ± 0.0 | 1.4 ± 0.5 |
| p-value | 0.72 | **<0.001** | 0.31 | 0.30 | 0.089 | 0.4884 | 0.7075 |

All values are mean ± SD; time points are with respect to injury; \*unpaired t-test (significant values are bolded); \*\*Kruskal-Wallis test.

### Injury Survival

For a full description of the typical response to this injury, including video, please refer to a previous publication.^19^ The survival of the rapid *vs*. slow infusion groups was 58% (7/12 subjects) *vs*. 67% (8/12 subjects), respectively; p >0.05 (Fisher exact test; see Kaplan-Meier plot in Fig. 2). The five subjects in the rapid group that expired prior to the end of the prescribed observation period were pronounced dead at 15, 26, 38, 40, and 43 min after injury; the four subjects in the slow group that similarly expired died at 30, 34, 35, and 53 min after injury. All deaths occurring prior to the prescribed observation period were preceded by terminal hypotension (i.e., no dysrhythmia, ventilatory inadequacy, or other contributing factors were noted).

**Fig. 2.**
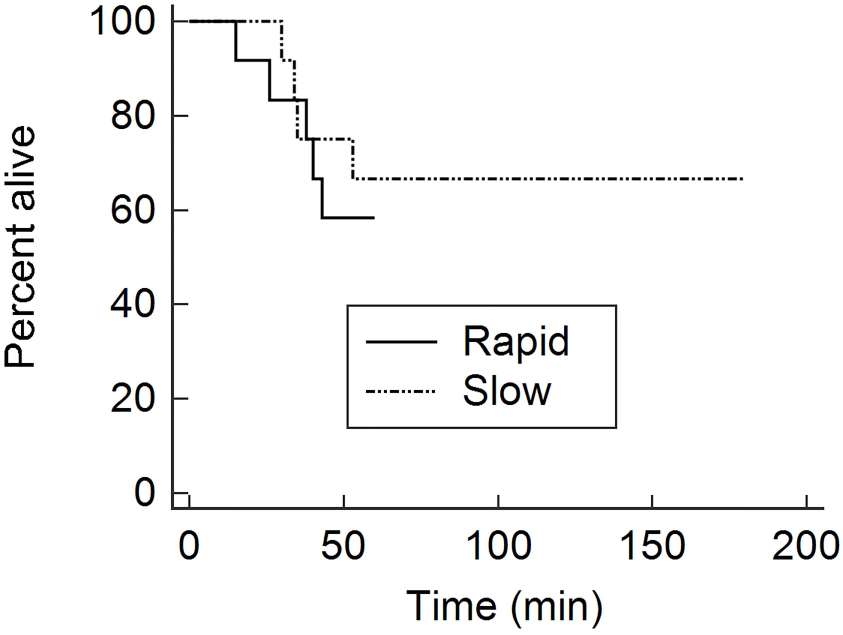
Kaplan Meier plot of injured porcine subjects treated with rapid *vs.* slow crystalloid resuscitation.

### Vital Sign Data

The MAP dropped precipitously (from >100 mm Hg to 30-60 mm Hg) within each group by 1-2 min after injury, but there were no significant differences in post-injury MAP between the rapid and slow groups (Table 2) at the 15 min or final time points. Interestingly, the MAP in the rapid group trended higher (nonsignificant) at the 15 min time point with respect to the slow group, but the opposite nonsignificant trend was present at the final time point (i.e., the MAP of the slow group trended higher). Although there was a small significant difference (~1°C) in temperature at the final time point, there were no post-injury differences between the two groups for heart rate and temperature (Table 2).

**Table 2.** Vital sign data.

| Infusion Rate | MAP (mm Hg) |  |  | Heart rate (beats/min) |  |  | Temperature (°C) |  |  |
| --- | --- | --- | --- | --- | --- | --- | --- | --- | --- |
|  | Initial | 15 min | Final | Initial | 15 min | Final | Initial | 15 min | Final |
| Rapid | 113 ± 14 | 55 ± 31 | 27 ± 16 | 107 ± 21 | 108 ± 22 | 100 ± 23 | 36.9 ± 0.6 | 36.5 ± 0.5 | 36.0 ± 0.6 |
| Slow | 105 ± 18 | 33 ± 17 | 39 ± 31 | 90 ± 19 | 110 ± 25 | 112 ± 23 | 36.3 ± 1.1 | 35.9 ± 1.0 | 35.1 ± 1.0 |
| *p-value | 0.52 | 0.11 | 0.48 | 0.09 | 0.99 | 0.69 | 0.23 | 0.15 | **0.010** |

MAP = Mean Arterial Pressure. All values are mean ± SD; time points are with respect to injury; \*Kruskal-Wallis test; significant tests are bolded.

### Blood Loss and Hematology Data

Post-injury blood loss was ~70% greater in the rapid group compared to the slow group (~2.7 vs. 1.6 L, respectively), but total post-injury crystalloid administered was not different (Table 1). The latter result was intentional per the experimental design; i.e., the total maximum crystalloid volume was set at 100 mL/kg for a subject in either group. Post-injury serum hemoglobin in the rapid group was about half the value of the slow group at both the 15 min and final time points; platelet concentration was lower in the rapid group at the 15 min time point only (Table 3). The QFA, PT, and INR were all more negatively affected in the rapid group compared to the slow group at the 15 time point (Table 4). Data on APTT were not analyzed secondary to an excessive number of missing values.

**Table 3.** Hematologic testing results.

| Infusion Rate | Hemoglobin (g/dL) | | | Platelets (x 1,000/μL) | | |
| --- | --- | --- | --- | --- | --- | --- |
|  | 0 min | 15 min | Final | 0 min | 15 min | Final |
| Rapid | 12.2 ± 0.9 | 4.6 ± 2.0 | 3.4 ± 2.3 | 306 ± 77 | 147 ± 66 | 122 ± 66 |
| Slow | 11.7 ± 0.9 | 8.4 ± 1.2 | 6.0 ± 2.2 | 315 ± 87 | 215 ± 55 | 171 ± 71 |
| p-value* | 0.39 | **<0.001** | **0.025** | 0.99 | **0.029** | 0.23 |

All values are mean ± SD; time points are with respect to injury; \*unpaired t-test; significant values are bolded.

**Table 4.**
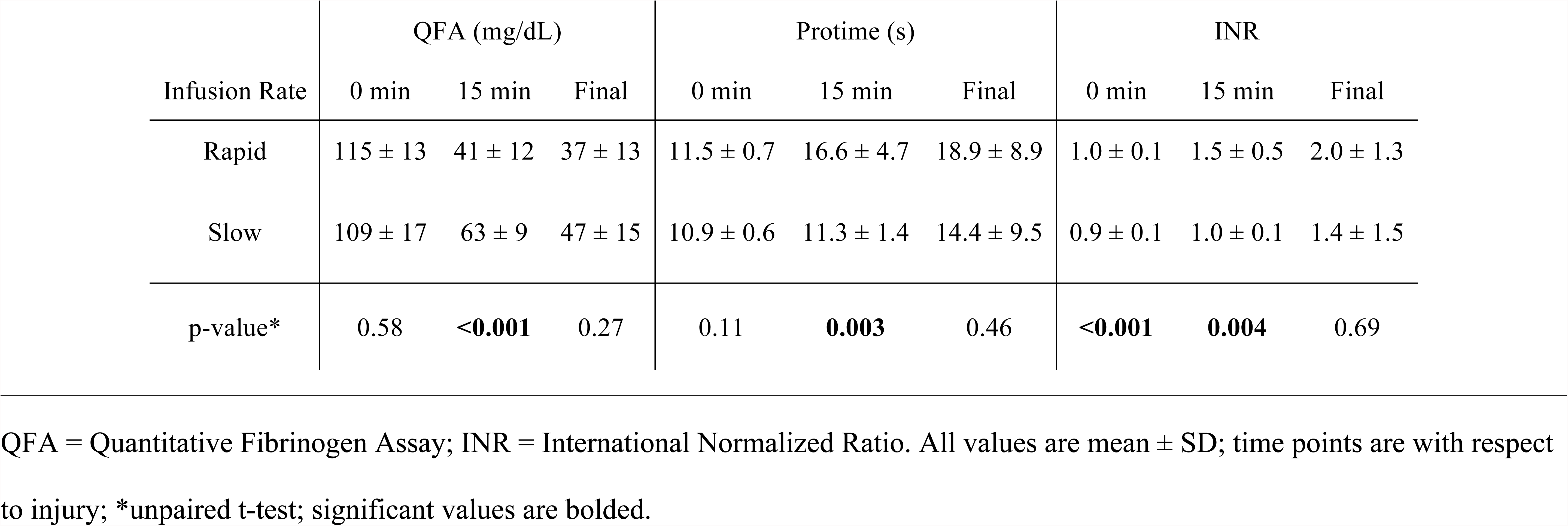
Coagulation testing results.

### Blood Gas and Necropsy Results

The within-group pH, pCO_2_, bicarbonate, and base excess all trended down after injury, but there were no differences in these values comparing the rapid vs. slow groups at any time point (see Statistical Analysis in the Supplemental Digital Content). End-tidal pCO_2_ was monitored but the electronic readout was not reliably captured, so comparative analysis of these data were not possible. Liver mass at necropsy not different between the groups (Table 1). Postmortem dissection of the explanted liver demonstrated that there was no difference between the two groups with respect to the number of hepatic or portal veins that were lacerated (Table 1).

## DISCUSSION

In this study of uncontrolled intraabdominal hemorrhage from a combined portohepatic venous injury in swine, it was found that for a fixed volume of resuscitation fluid, a 20 mL/min intravenous infusion rate (“slow”) resulted in a reduction in blood loss and an improvement in hemoglobin, platelet, and coagulation parameters compared to an infusion rate of 150 mL/min (“rapid”). The actual volume of post-injury crystalloid given did not differ between the two groups (this volume was capped at 100 mL/kg). So what was really tested in this study was the effect of the pump rate for a fixed volume of fluid. Essentially, subjects with the slower pump rate did better.

Countering the improved endpoints in the slow group was the observation that the final mean temperature was about 1°C greater in the rapid infusion group. This result likely was secondary to the fact that rapid group received the fixed volume of pre-warmed LR resuscitation fluid over a shorter time period. It is not clear whether the difference in final rectal temperature (36 vs. 35°C) was biologically relevant.

The subjects of this study were not randomized, because this study was not a pre-planned trial in the authors’ laboratory. Our primary task had been to develop hemostatic technologies for uncontrollable intraabdominal hemorrhage (unpublished data) using a porcine model,^19^ in which we utilized a rapid fluid-resuscitation protocol frequently used in these models.^20,25,29-31^ Since this protocol was not completely consistent with the TCCC Guidelines,^5^ and because we believed that rapid fluid resuscitation was negatively affecting our outcomes, we switched to a slower fluid resuscitation rate, but using the same total volume limit. Our subsequent impression was that the pigs were doing better with the slower rate. At that point we decided to perform a comparison of the fast vs. slow resuscitation groups, which thus produced the case-control design of this manuscript.

This study design is a flaw of this manuscript; the study groups are not strictly comparable. For example, the slow resuscitation protocol required extending the post-injury observation period to 180 min (compared to the 60 min observation period of the fast group), so that the total resuscitation volume in the fast *vs.* slow groups would be equal. However, despite the slow group having three times the duration of post-injury observation, this group still lost ~1 L less blood than the rapid group. We believe that this is a significant result that is not negated by the study design.

We could have compared the two resuscitation protocols at one hour, thus eliminating the issue of differing durations of post-injury monitoring, but then the total fluid volume would not have been equal between groups. Alternatively, we could have redone the study in a randomized fashion, using a 3 h observation period in all subjects. But given the convincing outcomes obtained with the case-control subjects reported herein, the involved regulatory committees would not have permitted these experiments to be redone. Another potential issue with this study is the lack of a no-treatment control group, which has been utilized in other preclinical studies of fluid resuscitation in hemorrhagic shock.^6,9,32,33^ Given the fact that the current TCCC guidelines^5^ do not recommend non-resuscitation in the prehospital phase, we decided not to include a no-treatment group in our study.

With 12 subjects per group, this study was not powered to detect a difference in survival between the rapid vs. slow crystalloid infusion groups. If the survival of the rapid infusion group is held constant at 58%, then in order to have 80% power to detect a significant difference (p < 0.05) with the Fisher exact test, the survival of the slow infusion group would need to be 100% with 17 subjects per group. Alternatively, in order to detect a survival difference of 20% at 80% power, the study would have required 85 subjects per group. So with only 12 subjects per group, the ability of the present study to detect a difference in survival was limited.

In addition to the above issues, other limitations of this study include no long-term survival data, data acquisition only under general anesthesia, and no data on the systemic inflammatory response. Moreover, there are controversial several aspects of porcine modeling of severe hemorrhage (including uncontrolled intraabdominal hemorrhage) which were not intended targets in this study, including the necessity of routine pre-injury splenectomy,^34,35^ the type of resuscitation fluid to employ,^33,36^ and the relative advantages of open vs. closed injury mechanisms.^19,20,25,28,37^ We utilized routine pre-injury splenectomy in this study based on the published experience of multiple groups in this field;^6,10,23-26^ we do acknowledge, however, that splenectomy for this purpose is controversial.^34,35^

In previous studies, 2-h survival in non-resuscitated pigs with uncontrolled aortic hemorrhage was 100%, while it was <50% in subjects resuscitated with bolused hypertonic saline/dextran (4 mL/kg over 1 min) or rapidly infused lactated Ringer’s (80 mL/kg over 9 min).^9^ Pigs undergoing a 40% blood volume bleed through a carotid arterial catheter and not resuscitated had somewhat improved coagulation parameters after a 2-h observation period compared to pigs who were fluid resuscitated.^33^ Pigs undergoing a grade V liver injury and not resuscitated shed less blood compared to fluid resuscitated pigs after a 2-h observation period,^32^ though blood pressure was marginally less in the former pigs. In a combined femoral/aortic hemorrhage model in pigs, resuscitation to a MAP of 40 mm Hg produced better 1-h survival than a resuscitation target of 80 mm Hg or no resuscitation at all.^6^

Regarding previous preclinical studies which specifically evaluated the effect of fluid administration rate on hemorrhagic shock, less data is available. Sondeen *et al.* found that the rate of the lactated Ringer’s resuscitation (100 *vs.* 300 mL/min) did not affect the time at which pigs rebled from an aortotomy;^21^ the major determinant of rebleeding (their primary endpoint) was a MAP greater than ~60 mm Hg. Pigs resuscitated with the 300 mL/min rate in this study, however, had a lower serum hemoglobin at the point of rebleeding. In regards to clinical studies, reports since the 1990’s have suggested^3, 7,11-13^ that if blood products are not available, then fluid restriction (i.e., hypotensive resuscitation) in the pre-hospital management of hemorrhagic shock may produce better outcomes.

Overall, the preclinical and clinical data support the use of low-volume resuscitation for prehospital management of hemorrhagic shock. The present study supports using a slow infusion rate as opposed to rapid infusion for a relatively large (100 mL/kg) fixed volume of crystalloid resuscitation. Whether additional benefit might be gained by combining a slow infusion rate with a low fixed volume of resuscitation fluid will require additional study.

A ready explanation for the apparent detrimental effect of rapid intravenous resuscitation for hemorrhagic shock seen in this study was hemodilution. Although both groups received the same fluid volume of crystalloid in the post-injury period, perhaps the slower infusion rate provided adequate time for that infused fluid to equilibrate into the extravascular space without causing hemodilution, while the rapid infusion rate overwhelmed the vascular space and diluted its contents. Furthermore, it has been demonstrated in animal models that saline infusion can activate systemic inflammation;^38, 39^ what role this phenomenon may have played in the present study was not determined.

Parameters describing a fluid maximum or fluid administration rate for prehospital crystalloid administration are not described in the current TCCC guidelines.^5^ While this study cannot provide these parameters, the data contained in this report can contribute to the ongoing discussion on how this aspect of prehospital management might evolve. In particular, it may ultimately be determined that a relatively slow intravenous infusion rate (as opposed to bolus infusion described in the current TCCC guidelines^5^) may be preferred for prehospital fluid resuscitation of the patient with hemorrhagic shock for which blood products are not immediately available.

### Conclusions

In a porcine model of uncontrolled intraabdominal hemorrhage, a relatively slow rate (20 mL/min) of intravenous crystalloid resuscitation produced improved short-term outcomes (blood loss, hemoglobin level, and coagulation parameters) compared to a more rapid rate (150 mL/min). The study was not powered to determine if there was a difference in short-term survival.

## Funding

This work was supported by a grant from the United States Army (W81XWH-11-1-0836).

## Conflicts of Interest

The authors report no conflicts of interest. The authors alone are responsible for the content and writing of the paper.

## Acknowledgements

This study is the result of work supported in part with resources and the use of facilities at the VA Nebraska-Western Iowa Health Care System. The authors would like to acknowledge the technical assistance of Chris Hansen, Dean Heimann, and Gerri Siford. Portions of this study were presented at the Annual Meeting of the Central Surgical Association (March 5-7, 2015 in Chicago, IL) and at the Military Health Systems Research Symposium (August 17-20, 2015 in Ft. Lauderdale, FL).

List of Supplemental Digital Content

1. Supplementary Table S1 (ARRIVE); pdf file
2. Statistical Analysis
3. Raw Data

(1 & 2 are combined into a single pdf file; 3 is an xls file)

## References

1. Blackbourne LH, Czarnik J, Mabry R, et al. Decreasing killed in action and died of wounds rates in combat wounded. J Trauma. 2010 Jul;69 Suppl 1:S1–4. PubMed PMID: 20622601.

2. Eastridge BJ, Hardin M, Cantrell J, et al. Died of wounds on the battlefield: causation and implications for improving combat casualty care. J Trauma. 2011 Jul;71(1 Suppl):S4–8. PubMed PMID: 21795876.

3. Shrestha B, Holcomb JB, Camp EA, et al. Damage-control resuscitation increases successful nonoperative management rates and survival after severe blunt liver injury. J Trauma Acute Care Surg. 2015;78(2):336–41. PubMed PMID: 25757120.

4. Butler F, Holcomb J, Schreiber M, et al. Fluid Resuscitation for Hemorrhagic Shock in Tactical Combat Casualty Care: TCCC Guidelines Change 14-01-2 June 2014. J Spec Op Med. 2013;14(3):13–38. PubMed PMID: 25344706.

5. United States Army Institute of Surgical Research Joint Trauma System: Tactical Combat Casualty Care Guidelines (version 03 June 2015). Accessed December 1, 2015. URL: https://www.jsomonline.org/TCCC.html-00.

6. Kowalenko T, Stern S, Dronen S, Wang X. Improved outcome with hypotensive resuscitation of uncontrolled hemorrhagic shock in a swine model. J Trauma. 1992 Sep;33(3):349–53; discussion 61-2. PubMed PMID: 1404501.

7. Morrison CA, Carrick MM, Norman MA, et al. Hypotensive resuscitation strategy reduces transfusion requirements and severe postoperative coagulopathy in trauma patients with hemorrhagic shock: preliminary results of a randomized controlled trial. J Trauma. 2011 Mar;70(3):652–63. PubMed PMID: 21610356.

8. Cannon W, Fraser J, Cowell E. The preventive treatment of wound shock. JAMA. 1918;70(9):618–21.

9. Bickell WH, Bruttig SP, Millnamow GA, et al. Use of hypertonic saline/dextran versus lactated Ringer’s solution as a resuscitation fluid after uncontrolled aortic hemorrhage in anesthetized swine. Ann Emerg Med. 1992 Sep;21(9):1077–85. PubMed PMID: 1381159.

10. Hildebrand F, Andruszkow H, Huber-Lang M, et al. Combined hemorrhage/trauma models in pigs-current state and future perspectives. Shock. 2013 Oct;40(4):247–73. PubMed PMID: 23856921.

11. Bickell WH, Wall MJ, Jr., Pepe PE, et al. Immediate versus delayed fluid resuscitation for hypotensive patients with penetrating torso injuries. N Engl J Med. 1994 Oct 27;331(17): 1105–9. PubMed PMID: 7935634.

12. Langan NR, Eckert M, Martin MJ. Changing Patterns of In-Hospital Deaths Following Implementation of Damage Control Resuscitation Practices in US Forward Military Treatment Facilities. JAMA surgery. 2014;149(9):904–12. PubMed PMID: 25029432.

13. Schreiber MA, Meier EN, Tisherman SA, et al. A controlled resuscitation strategy is feasible and safe in hypotensive trauma patients: Results of a prospective randomized pilot trial. J Trauma Acute Care Surg. 2015 Apr;78(4):687–97. PubMed PMID: 25807399. Pubmed Central PMCID: 4375962.

14. Kilkenny C, Browne WJ, Cuthill IC, et al. Improving bioscience research reporting: the ARRIVE guidelines for reporting animal research. PLoS Biol. 2010;8(6):e1000412. PubMed PMID: 20613859. Pubmed Central PMCID: PMC2893951.

15. Committee for the Update of the Guide for the Care and Use of Laboratory Animals. Guide for the Care and Use of Laboratory Animals: Washington, DC: The National Academies Press; 2011.

16. American Veterinary Medical Association Panel on Euthanasia. AVMA Guidelines for the Euthanasia of Animals: 2013 Edition. Schaumberg, IL: American Veterinary Medical Association; 2013.

17. Neter J, Wasserman W, Kutner MH. Applied Linear Statistical Models. 3rd ed. Boston: Irwin Publishing Co.; 1990.

18. Carlson MA, Calcaterra J, Johanning JM, et al. A totally recombinant human fibrin sealant. J Surg Res. 2014 Mar;187(1):334–42. PubMed PMID: 24169144. Epub 2013/10/31. eng.

19. Yanala UR, Johanning JM, Pipinos, II, et al. Development of a fatal noncompressible truncal hemorrhage model with combined hepatic and portal venous injury in normothermic normovolemic swine. PLoS One. 2014;9(9):e108293. PubMed PMID: 25251401. Pubmed Central PMCID: 4176969.

20. Duggan MJ, Mejaddam AY, Beagle J, et al. Development of a lethal, closed-abdomen grade V hepato-portal injury model in non-coagulopathic swine. J Surg Res. 2013 Jun 1;182(1):101–7. PubMed PMID: 22921917. Epub 2012/08/28. eng.

21. Sondeen JL, Coppes VG, Holcomb JB. Blood pressure at which rebleeding occurs after resuscitation in swine with aortic injury. J Trauma. 2003 May;54(5 Suppl):S110–7. PubMed PMID: 12768112.

22. Soller B, Zou F, Prince MD, et al. Comparison of Noninvasive pH and Blood Lactate as Predictors of Mortality in a Swine Hemorrhagic Shock with Restricted Volume Resuscitation Model. Shock. 2015 Aug;44 Suppl 1:90–5. PubMed PMID: 25526374. Pubmed Central PMCID: PMC4498648.

23. Sena MJ, Douglas G, Gerlach T, et al. A pilot study of the use of kaolin-impregnated gauze (Combat Gauze) for packing high-grade hepatic injuries in a hypothermic coagulopathic swine model. J Surg Res. 2013 Aug;183(2):704–9. PubMed PMID: 23541814.

24. Grottke O, Braunschweig T, Daheim N, et al. Effect of TachoSil in a coagulopathic pig model with blunt liver injuries. J Surg Res. 2011 Nov;171(1):234–9. PubMed PMID: 20452609.

25. Holcomb JB, Pusateri AE, Harris RA, et al. Effect of dry fibrin sealant dressings versus gauze packing on blood loss in grade V liver injuries in resuscitated swine. J Trauma. 1999;46(1):49–57. PubMed PMID: 1999129463. English.

26. Schreiber MA, Holcomb JB, Hedner U, et al. The effect of recombinant factor VIIa on coagulopathic pigs with grade V liver injuries. J Trauma. 2002 Aug;53(2):252–7; discussion 7-9. PubMed PMID: 12169930.

27. Pusateri AE, Modrow HE, Harris RA, et al. Advanced hemostatic dressing development program: animal model selection criteria and results of a study of nine hemostatic dressings in a model of severe large venous hemorrhage and hepatic injury in Swine. J Trauma. 2003 Sep;55(3):518–26. PubMed PMID: 14501897.

28. Duggan MJ, Rago A, Marini J, et al. Development of a lethal, closed-abdomen, arterial hemorrhage model in noncoagulopathic swine. J Surg Res. 2014 Apr;187(2):536–41. PubMed PMID: 24398305.

29. Duggan M, Rago A, Sharma U, et al. Self-expanding polyurethane polymer improves survival in a model of noncompressible massive abdominal hemorrhage. J Trauma Acute Care Surg. 2013 Jun;74(6): 1462–7. PubMed PMID: 23694873. Epub 2013/05/23. eng.

30. Delgado AV, Kheirabadi BS, Fruchterman TM, et al. A novel biologic hemostatic dressing (fibrin patch) reduces blood loss and resuscitation volume and improves survival in hypothermic, coagulopathic Swine with grade V liver injury. J Trauma. 2008 Jan;64(1):75–80. PubMed PMID: 18188102.

31. Kiraly LN, Differding JA, Enomoto TM, et al. Resuscitation with normal saline (NS) vs. lactated ringers (LR) modulates hypercoagulability and leads to increased blood loss in an uncontrolled hemorrhagic shock swine model. J Trauma. 2006 Jul;61(1):57–64; discussion −5. PubMed PMID: 16832250.

32. Riha GM, Kunio NR, Van PY, et al. Hextend and 7.5% hypertonic saline with Dextran are equivalent to Lactated Ringer’s in a swine model of initial resuscitation of uncontrolled hemorrhagic shock. J Trauma. 2011 Dec;71(6):1755–60. PubMed PMID: 22182885.

33. Via D, Kaufmann C, Anderson D, et al. Effect of hydroxyethyl starch on coagulopathy in a swine model of hemorrhagic shock resuscitation. J Trauma. 2001 Jun;50(6): 1076–82. PubMed PMID: 11426123.

34. Bebarta VS, Daheshia M, Ross JD. The significance of splenectomy in experimental swine models of controlled hemorrhagic shock. J Trauma Acute Care Surg. 2013;75(5):920. PubMed PMID: 24158218.

35. Boysen SR, Caulkett NA, Brookfield CE, et al. Splenectomy Versus Sham Splenectomy in a Swine Model of Controlled Hemorrhagic Shock. Shock. 2016 Oct;46(4):439–46. PubMed PMID: 26974424.

36. Sillesen M, Johansson PI, Rasmussen LS, et al. Fresh frozen plasma resuscitation attenuates platelet dysfunction compared with normal saline in a large animal model of multisystem trauma. J Trauma Acute Care Surg. 2014 Apr;76(4):998–1007. PubMed PMID: 24662863.

37. Cho SD, Holcomb JB, Tieu BH, et al. Reproducibility of an animal model simulating complex combat-related injury in a multiple-institution format. Shock. 2009 Jan;31(1):87–96. PubMed PMID: 18497710.

38. Kellum JA, Song M, Almasri E. Hyperchloremic acidosis increases circulating inflammatory molecules in experimental sepsis. Chest. 2006 Oct;130(4):962–7. PubMed PMID: 17035425.

39. Aksu U, Bezemer R, Yavuz B, et al. Balanced vs unbalanced crystalloid resuscitation in a near-fatal model of hemorrhagic shock and the effects on renal oxygenation, oxidative stress, and inflammation. Resuscitation. 2012 Jun;83(6):767–73. PubMed PMID: 22142654.

